# A coarse-grained elastic network atom contact model and its use in the simulation of protein dynamics and the prediction of the effect of mutations

**DOI:** 10.1101/001495

**Authors:** Vincent Frappier, Rafael Najmanovich

## Abstract

Normal mode analysis (NMA) methods are widely used to study dynamic aspects of protein structures. Two critical components of NMA methods are coarse-graining in the level of simplification used to represent protein structures and the choice of potential energy functional form. There is a trade-off between speed and accuracy in different choices. In one extreme one finds accurate but slow molecular-dynamics based methods with all-atom representations and detailed atom potentials. On the other extreme, fast elastic network model (ENM) methods with C_α_-only representations and simplified potentials that based on geometry alone, thus oblivious to protein sequence. Here we present ENCoM, an Elastic Network Contact Model that employs a potential energy function that includes a pairwise atom-type non-bonded interaction term and thus makes it possible to consider the effect of the specific nature of amino-acids on dynamics within the context of NMA. ENCoM is as fast as existing ENM methods and outperforms such methods in the generation of conformational ensembles. Here we introduce a new application for NMA methods with the use of ENCoM in the prediction of the effect of mutations on protein stability. While existing methods are based on machine learning or enthalpic considerations, the use of ENCoM, based on vibrational normal modes, is based on entropic considerations. This represents a novel area of application for NMA methods and a novel approach for the prediction of the effect of mutations. We compare ENCoM to a large number of methods in terms of accuracy and self-consistency. We show that the accuracy of ENCoM is comparable to that of the best existing methods. We show that existing methods are biased towards the prediction of destabilizing mutations and that ENCoM is less biased at predicting stabilizing mutations.

**Author Summary:** Normal mode analysis (NMA) methods can be used to explore the ensemble of potential movements around an equilibrium conformation by mean of calculating the eigenvectors and eigenvalues associated to different normal modes. Each normal mode represents one particular set of global collective, correlated and complex, form of motion of all atoms in the system. Any conformation around equilibrium can be represented as a linear combination of amplitudes associated to each normal mode. Differences in the magnitudes of the set of eigenvalues between two structures can be used to calculate differences in entropy. Coarse-grained NMA methods utilize a simplified potential and representation of the protein structure and thus decrease the computational time necessary to calculate eigenvectors and their respective eigenvalues. Here we present ENCoM the first coarse-grained NMA method to consider side-chain atomic interactions and thus able to calculate the effect of mutations on eigenvectors and eigenvalues. Such differences in turn are related to entropic differences and can thus be used to predict the effect of mutations on protein stability. ENCoM performs better than existing NMA methods with respect to different traditional applications of NMA methods (such as conformational sampling) or comparably for the prediction or crystallographic b-factors. ENCoM is the first NMA method that can be used to predict the effect of mutations on protein stability. Comparing ENCoM to a large set of dedicated methods for the prediction of the effect of mutations on protein stability shows that ENCoM performs better than existing methods considering a combination of prediction ability (particularly on stabilizing mutations) and bias (how does the prediction of a forward or back mutations differ). ENCoM is the first entropy-based method developed to predict the effect of mutations on protein stability.

## INTRODUCTION

Biological macromolecules are dynamic objects. In the case of proteins, such movements form a continuum ranging from bond and angle vibrations, sub-rotameric and rotameric side-chain rearrangements [1], loop or domain movements through to folding. Such movements are closely related to function playing important roles in most processes such as enzyme catalysis [2], signal transduction [3] and molecular recognition [4] among others. While the number of proteins with known structure is vast with around 85 K structures for over 35 K protein chains (at 90% sequence identity) in the PDB database [5], our view of protein structure tends to be somewhat biased, even if unconsciously, towards considering such macromolecules as rigid objects. This is due in part to the static nature of images used in publications to guide our interpretations of how structural details influence protein function. However, the main reason is that most known structures were solved using X-ray crystallography [6] where dynamic proprieties are limited to b-factors and the observation of alternative locations. Despite this, it is common to analyze larger conformational changes using X-ray structures with the comparison of different crystal structures for the same protein obtained in different conditions or bound to different partners (protein, ligand, nucleic acid). It is particularly necessary to consider the potential effect of crystal packing [7,8] when studying dynamic properties using X-ray structures.

Nuclear magnetic resonance (NMR) is a powerful technique that gives more direct information regarding protein dynamics [9,10]. Different NMR methodologies probe distinct timescales covering 15 orders of magnitude from 10^−12^ s side chain rotations via nuclear spin relaxation to 10^3^ s using real time NMR [10]. In practice, there is a limitation on the size of proteins that can be studied (between 50-100 kDa) although this boundary is being continuously pushed [11] providing at least partial dynamic information on extremely large systems [12]. However, only a small portion (around 10%) of the available proteins structures in the Protein Data Bank (PDB) are the result of NMR experiments [5].

Molecular dynamic simulations numerically solve the classical equations of motion for an ensemble of atoms whose interactions are modeled using empirical potential energy functions [13–15]. At each time step the positions and velocities of each atom are calculated based on their current position and velocity as a result of the forces exerted by the rest of the system. The first MD simulation of a protein (Bovine Pancreatic Trypsin Inhibitor, BPTI) ran for a total 8.8ps [16] followed by slightly longer simulations (up to 56 ps) [17]. A film of the latter can be seen online (http://youtu.be/_hMa6G0ZoPQ). Despite using simplified potentials and structure representations (implicit hydrogen atoms) as well as ignoring the solvent, these first simulations showed large oscillations around the equilibrium structure, concerted loop motions and hydrogen bond fluctuations that correlate with experimental observations. Nowadays, the latest breakthroughs in molecular dynamics simulations deal with biological processes that take place over longer timescales. For example, protein folding [18], transmembrane receptor activation [19,20] and ligand binding [21]. These simulations require substantial computer power or purpose built hardware such as Anton that pushes the current limit of MD simulations to the millisecond range [22]. Despite powerful freely available programs like NAMD [23] and GROMACS [24] and the raise of computational power over the last decade, longer simulations reaching timescales where most biological processes take place are still state-of-the-art.

Normal modes are long established in the analysis of the vibrational properties of molecules in physical chemistry [25]. Their application to the study of proteins dates back to just over 30 years [26–30]. These earlier Normal Mode Analysis (NMA) methods utilized either internal or Cartesian coordinates and complex potentials (at times the same ones used in MD). As with earlier MD methods their application was restricted to relatively small proteins. Size limitations notwithstanding, these early studies were sufficient to demonstrate the existence of modes representing concerted delocalized motions, showing a facet of protein dynamics that is difficult to access with MD methods. Some simplifications were later introduced and shown to have little effect on the slowest vibrational modes and their utility to predict certain molecular properties such as crystallographic b-factors. These simplifications included the use of a single-parameter potential [31], blocks of consecutive amino acids considered as units (nodes) [32] and the assumptions of isotropic [33] fluctuations in the Gaussian Network Model (GNM) or anisotropic fluctuations [34]. These approximations have drastically reduced the computational time required, thus permitting a much broader exploration of conformational space using conventional desktop computers in a matter of minutes. Of these, the most amply used method is the Anisotropic Network Model (ANM) [35,36]. ANM is often referred simply as an elastic network model; one should however bear in mind that all normal mode analysis methods are examples of elastic network models. ANM uses a simple Hook potential that connects every node (a point mass defined at the position of an alpha carbon), within a predetermined cut-off distance (usually 18Å). More recently, a simplified model, called Spring generalized Tensor Model (STeM), that uses a potential function with four terms (covalent bond stretching, angle bending, dihedral angle torsion and non-bonded interaction) has been proposed [37]. The normal mode analysis of a macromolecule produces a set of modes (eigenvectors and their respective eigenvalues) that represent possible movements. Any conformation of the macromolecule can in principle be reached from any other using a linear combination of amplitudes associated to eigenvectors.

It is essential however to not loose sight of the limitations of normal mode analysis methods. Namely, normal modes tell us absolutely nothing about the actual dynamics of a protein in the sense of the evolution in time of atomic coordinates. Plainly speaking, normal mode analysis is informative about the possible movements but not actual movements. Additionally, normal modes tell us of the possible movements around equilibrium. These two caveats clearly place normal mode analysis and molecular dynamics apart. First, molecular dynamics gives an actual dynamics (insofar as the potential is realistic and quantum effects can be ignored). Second, while the equilibrium state (or the starting conformation) affects the dynamics, one can explore biologically relevant timescales given sufficient computational resources to perform long simulations.

All coarse-grained NMA models only use the geometry of the protein backbone (via C_α_ Cartesian position) disregarding the nature of the corresponding amino acid, in doing so a lot of information is lost. By definition, such models cannot account for the effect of mutations on protein dynamics. It has been shown that different amino acids interact differently and that single mutations can have a high impact on protein function and stability [38–40]. Mutations on non-catalytic residues that participate into concerted (correlated) movements have been shown to disrupt protein function in NMR relaxation experiments [41–43]. Several cases have been documented of mutations that don’t affect the global fold of the protein, but affect protein dynamics and disrupt enzyme function [44].

To overcome this limitation of coarse-grained NMA methods while maintaining the advantages of simplified elastic network models, we developed a new mixed coarse-grained NMA model called Elastic Network Contact Model (ENCoM). ENCoM employs a potential function based on the four bodies potential of STeM with an addition to take in consideration the nature and the orientation of side chains. Side-chain atomic contacts are used to modulate the long range interaction term with a factor based on the surface area in contact [45] and the type of each atom in contact. Additionally, we introduce a non-specific version of ENCoM (ENCoM_ns_) where all interactions between atom types are the same. ENCoM and ENCoM_ns_ were validated trough comparison to ANM, GNM and STeM with respect to the prediction of crystallographic b-factors and conformational changes, two properties conventionally used to test ENM methods. Moreover, we test the ability of ENCoM and ENCoM_ns_ to predict the effect of mutations with respect to protein stability and compare the ability of ENCoM and ENCoM_ns_ to a large number of existing methods specifically designed for the prediction of the effect of single point mutations on protein stability. Finally, we use ENCoM to predict the effect of mutations on protein function in the absence of any effects on protein stability.

## RESULTS

### Correlation between experimental and predicted crystallographic b-factors

We utilized a dataset of 113 non-redundant high-resolution crystal structures [46] to predict b-factors using the calculated ENCoM eigenvectors and eigenvalues as described previously [35] (Equation 4). We compared the predicted b-factors using ENCoM, ENCoM_ns_, ANM, STeM and GNM to the experimental C_α_ b-factors for the above dataset (Supplementary Table I). For each protein we calculate the Pearson correlation between experimental and predicted values. The results in Figure 1 represent the bootstrapping average of 10000 iterations. We observe that while comparable, ENCoM (median = 0.54) and ENCoM_ns_ (median = 0.56) have lower median values than STeM (median = 0.60) and GNM (median = 0.59) but similar or higher than ANM (median = 0.54). It should be noted that it is possible to find specific parameter sets that maximize b-factor correlations beyond the values obtained with STeM and GNM (see methods). However we observe a trade-off between the prediction of b-factors on one side and overlap and the effect of mutations on the other (see methods). Ultimately we opted for a parameter set that maximizes overlap and the prediction of mutations with complete disregard to b-factor predictions. Nonetheless, as shown below (Figure 10), even the lower correlations obtained with ENCoM are sufficiently high to detect functionally relevant local variations in b-factors as a result of mutations. As GNM does not provide information on the direction of movements or the effect of mutations, it is not considered further in the present study.

**Figure 1.**
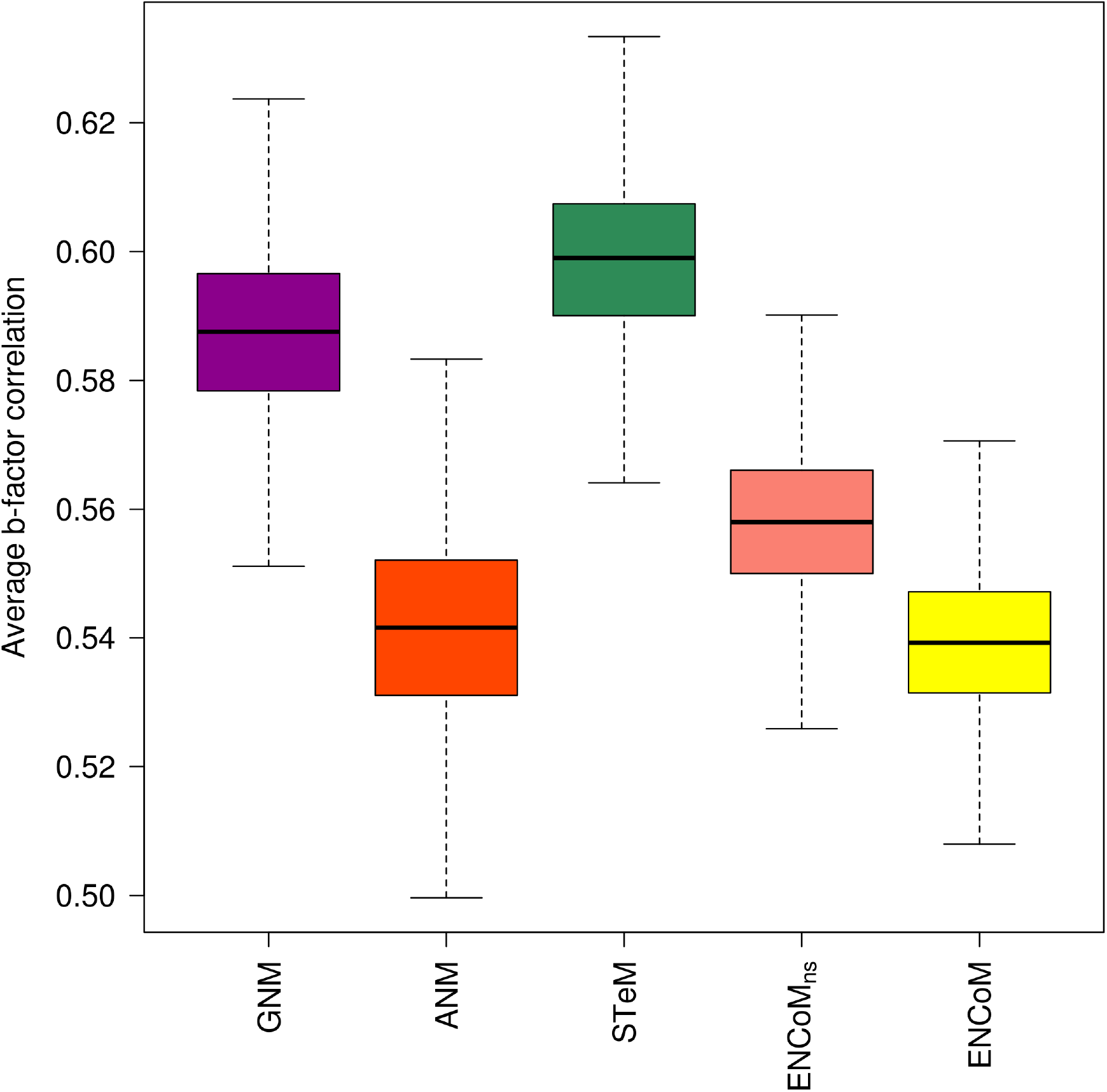
Correlation between predicted and experimental b-factors for different ENM models. Box plots represent the average correlations from a non-redundant dataset containing 113 proteins generated from 10000 resampling bootstrapping iterations. ANM and ENCoM have comparable correlations but lower than GNM and STeM.

### Exploration of conformational space

By definition, any conformation of a protein can be described as a linear combination of amplitudes associated to the eigenvectors representing normal modes. It should be stressed that such conformations are as precise as the choice of structure representation used and correct within the quadratic approximation of the potential around equilibrium. Those limitations notwithstanding, one application of NMA is to explore the conformational space of macromolecules using such linear combinations of amplitudes. Pairs of distinct protein conformations, often obtained by X-ray crystallography are used to assess the extent to which the eigenvectors calculated from a starting conformation could generate movements that could lead to conformational changes in the direction of a target conformation. Rather than an optimization to determine the amplitudes for a linear combination of eigenvectors, this is often simplified to the analysis of the overlap (Equation 5), i.e., the determination of the single largest contribution from a single eigenvector towards the target conformation. In a sense the overlap represents a lower bound on the ability to predict conformational changes without requiring the use of an optimization process.

The analysis of overlap for ANM, STeM, ENCoM and ENCoM_ns_ was performed using the Protein Structural Change Database (PSCDB) [47], which contains 839 pairs of protein structures undergoing conformational change upon ligand binding. The authors classify those changes into seven types: coupled domain motions (59 entries), independent domain motions (70 entries), coupled local motions (125 entries), independent local motions (135 entries), burying ligand motions (104 entries), no significant motion (311 entries) and other type of motions (35 entries). The independent movements are movements that don’t affect the binding pocket, while dependent movements are necessary to accommodate ligands in the pose found in the bound (holo) form. Burying movements are associated with a significant change of the solvent accessible surfaces of the ligand, but with small structural changes (backbone RMSD variation lower than 1Å). Despite differentiating between types of movements based on the ligands, the ligands were not used as part of the normal mode analysis. Since side-chain movements associated to the burying movements cannot be predicted with coarse-grained NMA methods, we restrict the analysis to domain and loop movements [48] as these involve backbone movements amenable to analysis using coarse grained NMA methods. For practical purposes, in order to simplify the calculations in this large-scale analysis, NMR structures were not considered. It is worth stressing however that all NMA methods presented here don’t have any restriction with respect to the structure determination method and can also be used with modeled structures. A total of 736 conformational changes, half representing apo to holo changes and the other half holo to apo (in total 368 entries from PSCDB) are used in this study (Supplementary Table II).

Overlap calculations were performed from the unbound (apo) form to the bound form (holo) and from the bound form to the unbound form. Bootstrapped results based onto the best overlap found within the first 10 slowest modes [49,50] for the different types of conformational changes, domain or loop are shown in figures 2 and 3 respectively. In each case a set of box-plots represent the performance of the four methods being compared, namely STeM, ANM, ENCoM and ENCoM_ns_. The left-most set of box-plots represents the average over all data while subsequent sets represent distinct subsets of the dataset as labeled. The first observation (comparing Figures 2 and 3) is that all tested NMA models show higher average overlaps for domain movements (Figure 2) than loop movements (Figure 3). This confirms earlier observations that NMA methods capture essential cooperative global (delocalized) movements associated with domain movements [48]. Loop movements on the other hand are likely to come about from a more fine tuned combination of normal mode amplitudes than what can be adequately described with a single eigenvector as measured by the overlap.

**Figure 2.**
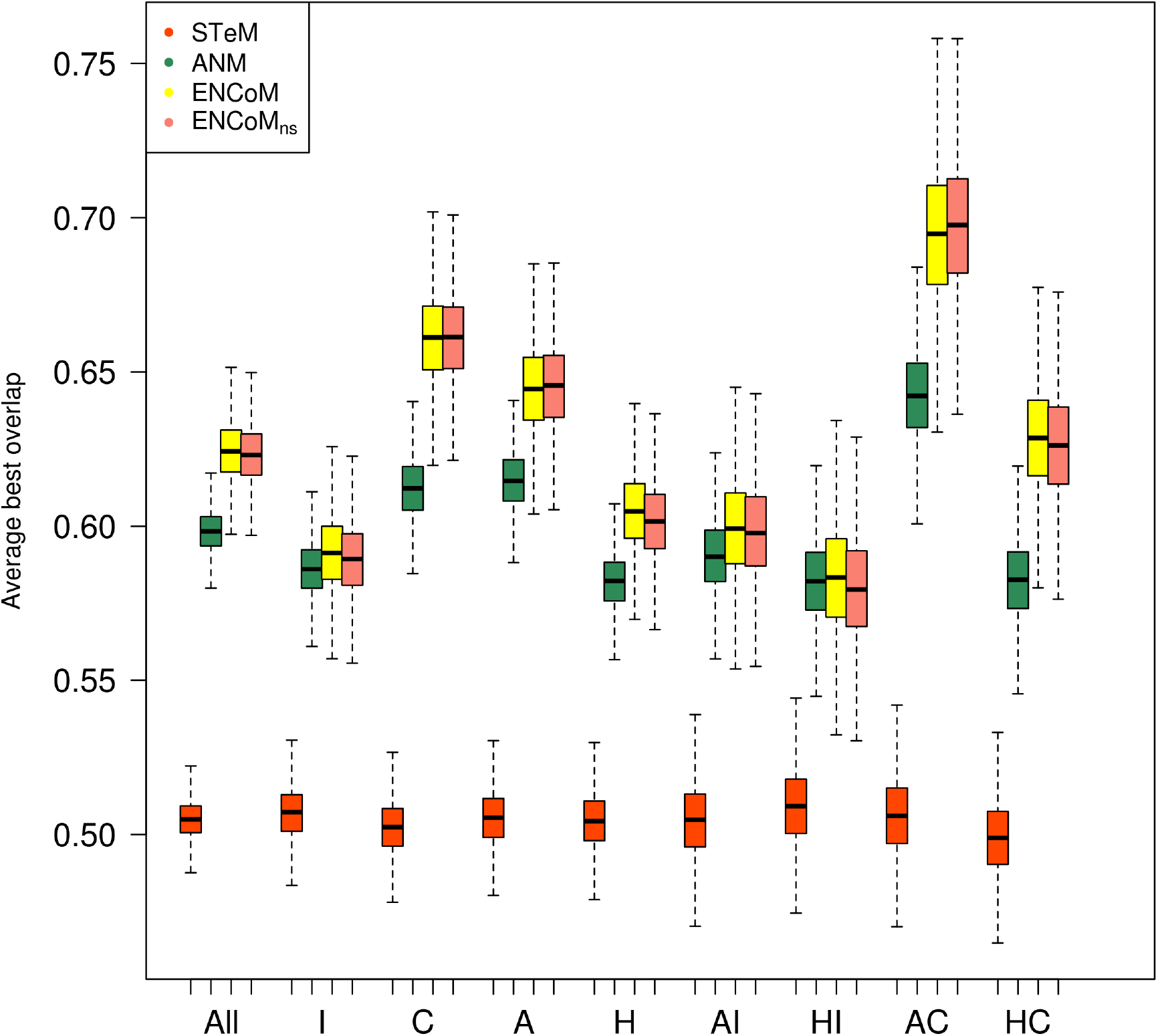
Prediction of domain motions. The best overlap found within the 10 slowest internal motion modes for different NMA models on domain movements. Legend: All (all cases, N = 248), I (Independent movements, N = 130), C (Coupled movements, N = 117), A (apo form, N = 124) and H (holo form, N = 124). AI (apo form independent, N = 130), HI (holo independent, N = 130), AC (apo coupled, N = 116) and HC (holo coupled, N = 116). Box plots generated from 10000 resampling bootstrapping iterations. ENCoM/ENCoM_ns_ outperform ANM and STeM on all types of motion.

**Figure 3.**
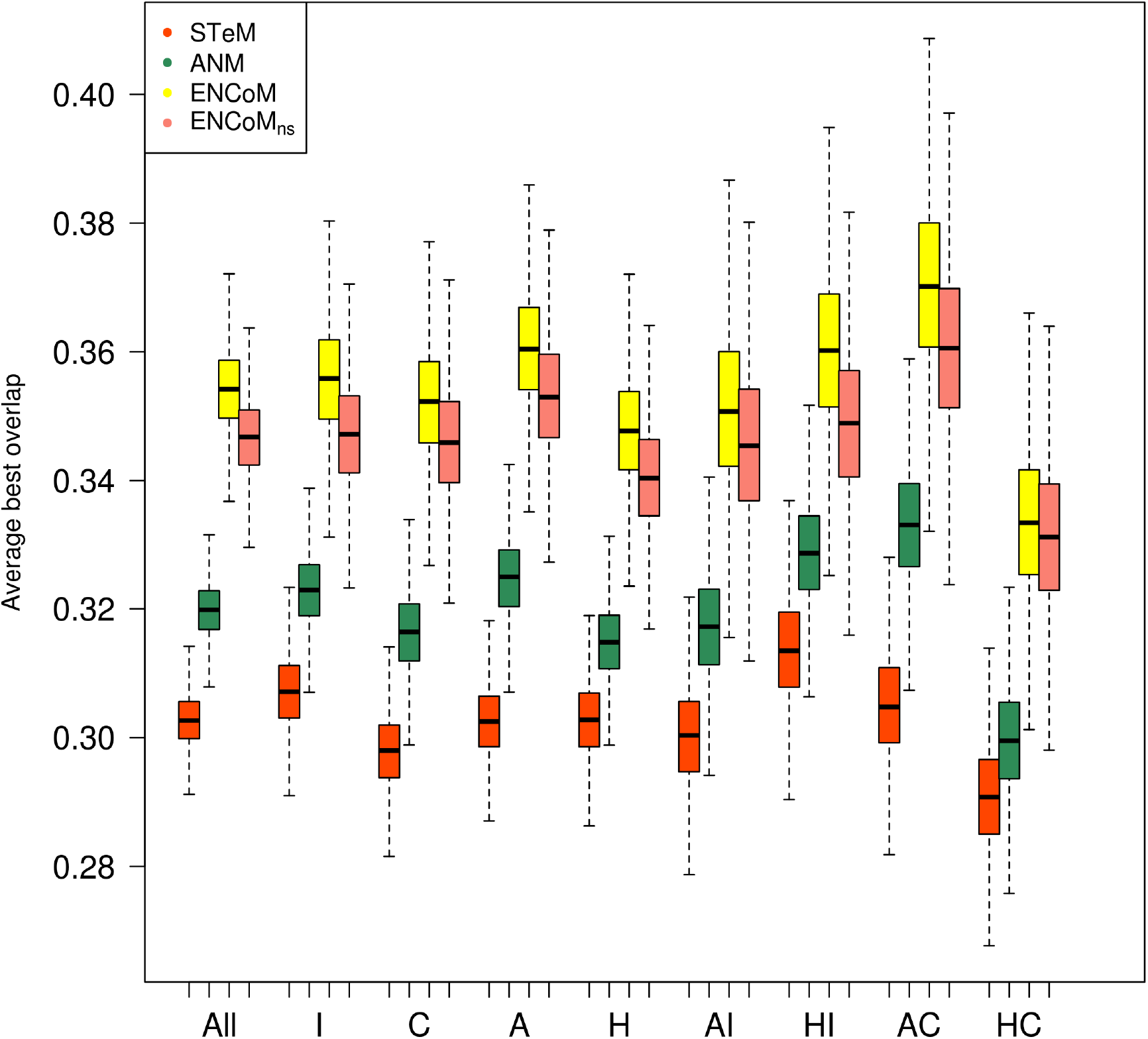
Prediction of loop conformational change. The best overlap found within the 10 slowest internal motion modes for different NMA models on loop movements. Acronyms are the same as in Figure 2. Number of cases: All (488), I (252), C (236), A (244) and H (244). AI (126), HI (126), AC (118) and HC (118). Box plots generated from 10000 resampling bootstrapping iterations. ENCoM/ENCoM_ns_ outperform ANM and STeM on all types of motion. The prediction of loop motions is much harder and here the difference between ENCoM and ENCoM_ns_ are more pronounced.

The second observation is that STeM performs quite poorly compared to other methods irrespective of the type of movement (domain or loop). This is somewhat surprising when one compares with ENCoM or ENCoM_ns_ considering how similar the potentials are. This suggests that the modulation of interactions by the surface area in contact (the *β*_*ij*_ terms in Equation 1) of the corresponding side-chains as well as the specific parameters used are crucial.

Focusing for a moment on domain movements (Figure 2), ENCoM/ENCoM_ns_ outperform all other methods for domain movements in general as well as for every sub category of types of motions therein. Independent movements show lower overlaps than coupled ones, a fraction of those movements may not be biologically relevant due to crystal packing. Interestingly, while there are no differences between the overlap for independent movements starting from the apo or holo forms, this is not the case for coupled movements. In this case (right-most two sets in Figure 2), it is easier to use the apo (unbound) form to predict the holo (bound) form, suggesting that the lower packing in the apo form (as this are frequently more open) generates eigenvectors that favor a more comprehensive exploration of conformational space.

Lastly, with respect to loop movements (Figure 3), while it is more difficult to obtain good overlaps irrespective of the method or type of structure used, overall ENCoM/ENCoM_ns_ again outperforms ANM. Some of the same patterns observed for domain movements are repeated here. For example, the higher overlap for coupled apo versus holo movements.

We observe that ENCoM_ns_ consistently performs almost as well as ENCoM irrespective of the type of motion used (all sets in Figures 2 and 3). As side-chain conformations in crystal structures tent to minimize unfavorable interactions, the modulation of interactions by atom types that differentiate ENCoM from ENCoM_ns_ plays as minor but still positive role.

### Prediction of the effect of mutations

Normal mode resonance frequencies (eigenvalues) are related to vibrational entropy [51,52] (see methods). Therefore, it is reasonable to assume that the information contained in the eigenvectors can be used to infer differences in protein stability between two structures differing by a mutation under the assumption that the mutation does not drastically affect the equilibrium structure. A mutation may affect stability due to an increasing in the entropy of the folded state by lowering its resonance frequencies, thus making more microstates accessible around the equilibrium.

We utilize experimental data from the ProTherm database [53] on the thermodynamic effect of mutations to validate the use of ENCoM to predict protein stability. Here we benefit from the manual curation efforts previously performed to generate a non-redundant subset of ProTherm comprising 303 mutations used for the validation of the PoPMuSiC-2.0 [54]. The dataset contains 45 stabilizing mutations (ΔΔG < −0.5 kcal/mol), 84 neutral mutations (ΔΔG [−0.5,0.5] kcal/mol) and 174 destabilizing mutations (ΔΔG > 0.5 kcal/mol) (Supplementary Table II). Each protein in the dataset have at least one structure in the PDB database [55]. As we calculate the eigenvectors in the mutated form we require model structures of the mutants. We generate such models using Modeller [56] and are thus assuming that the mutation does not drastically affect the structure.

In the present work we predict the effect of mutations (Equation 6) for ENCoM, ENCoM_ns_, ANM and STeM and compare the results to existing methods for the prediction of the effect of mutations using the PoPMuSiC-2.0 dataset above. We compare our results to those reported by Dehouck *et al.* [54] for different existing techniques: CUPSAT, a Boltzman mean-force potential [57]; DMutant, an atom-based distance potential [58]; PoPMuSiC-2.0, a neural network based method [54]; Eris, a force field based approach [59]; I-Mutant 2.0, a support vector machine method [60]; and AUTO-MUTE, a knowledge based four body potential [61]. We used the same dataset to generate the data for FoldX 3.0, an empirical full atom force-field [62] and Rosetta [63], based on the knowledge based Rosetta energy function. A negative control model was build with a randomized reshuffling of the experimental data. Figure 4 presents RMSE results for each model. The raw data for the 303 mutations is available in Supplementary Table III.

**Figure 4.**
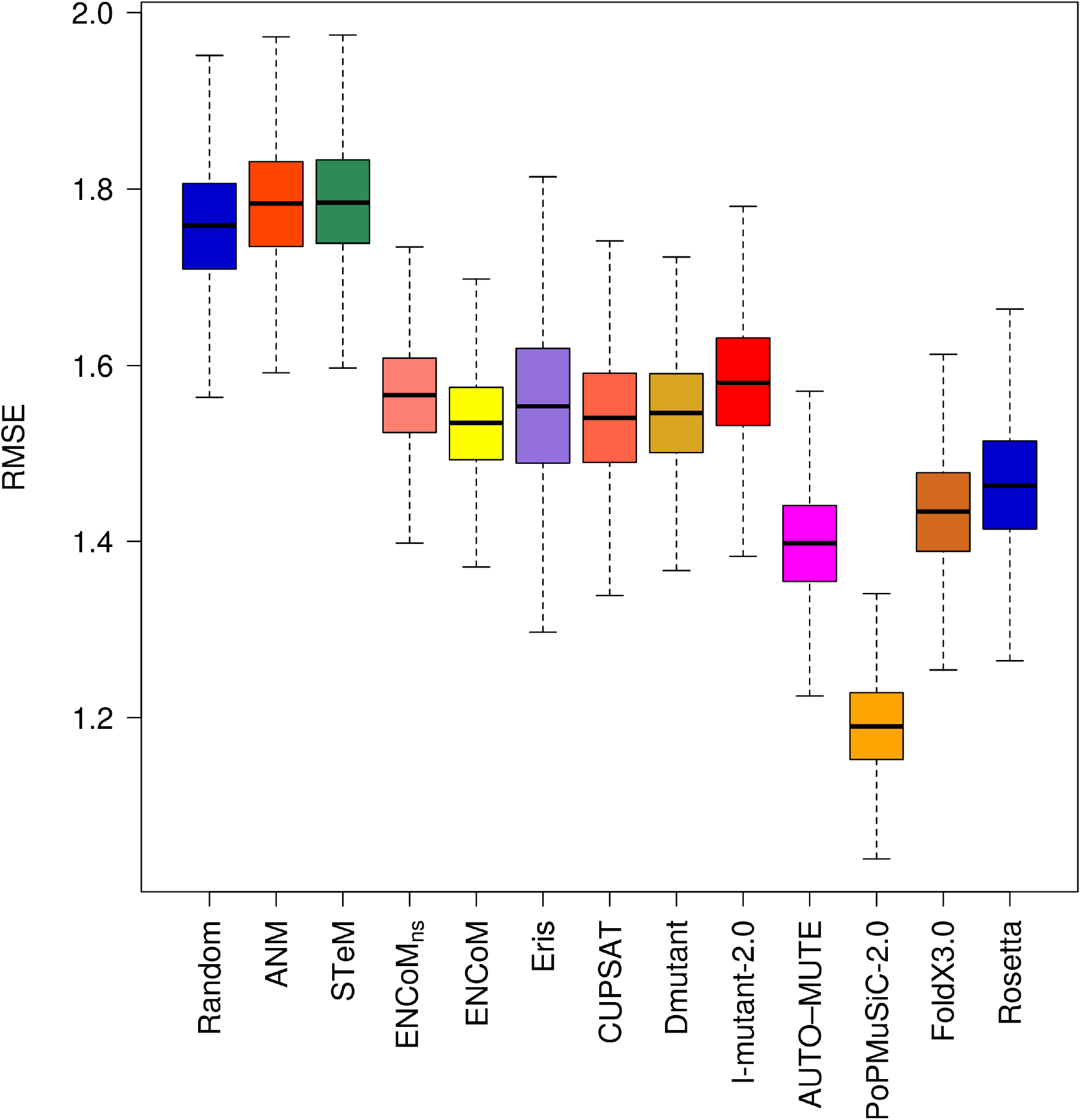
Root Mean Square Error (RMSE) on the prediction of the effect of mutations. RMSE of the linear regression through the origin between experimental and predicted variations in free energy variations (ΔΔG). Box plots generated from 10000 resampling bootstrapping iterations on the entire PoPMuSiC-2.0 dataset (N = 303). With the caveat that the dataset is biased towards destabilizing mutations, ENCoM/ENCoM_ns_ predict the effect of mutations with similar overall RMSE as most other methods.

ANM and STeM are as good as the random model when considering all types of mutations together (Figure 4). This is not surprising as the potentials used ANM and STeM are exclusively geometry-based and are thus agnostic to sequence. ENCoM_ns_, ENCoM, Eris, CUPSAT, DMutant, I-Mutant 2.0 and give similar results and predict significantly better than the random model. AUTO-MUTE, FoldX 3.0, Rosetta and in particular PoPMuSiC-2.0 outperform all of the other models.

The RMSE values for the subset of 174 destabilizing mutations (Figure 5) shows similar trends as the whole dataset with the exceptions of DMutant losing performance and PoPMuSiC-2.0 as well as AUTO-MUTE gaining performance compared to the others. It is important to stress that the low RMSE of PoPMuSiC-2.0 on the overall dataset is to a great extent due to its ability to predict destabilizing mutations.

**Figure 5.**
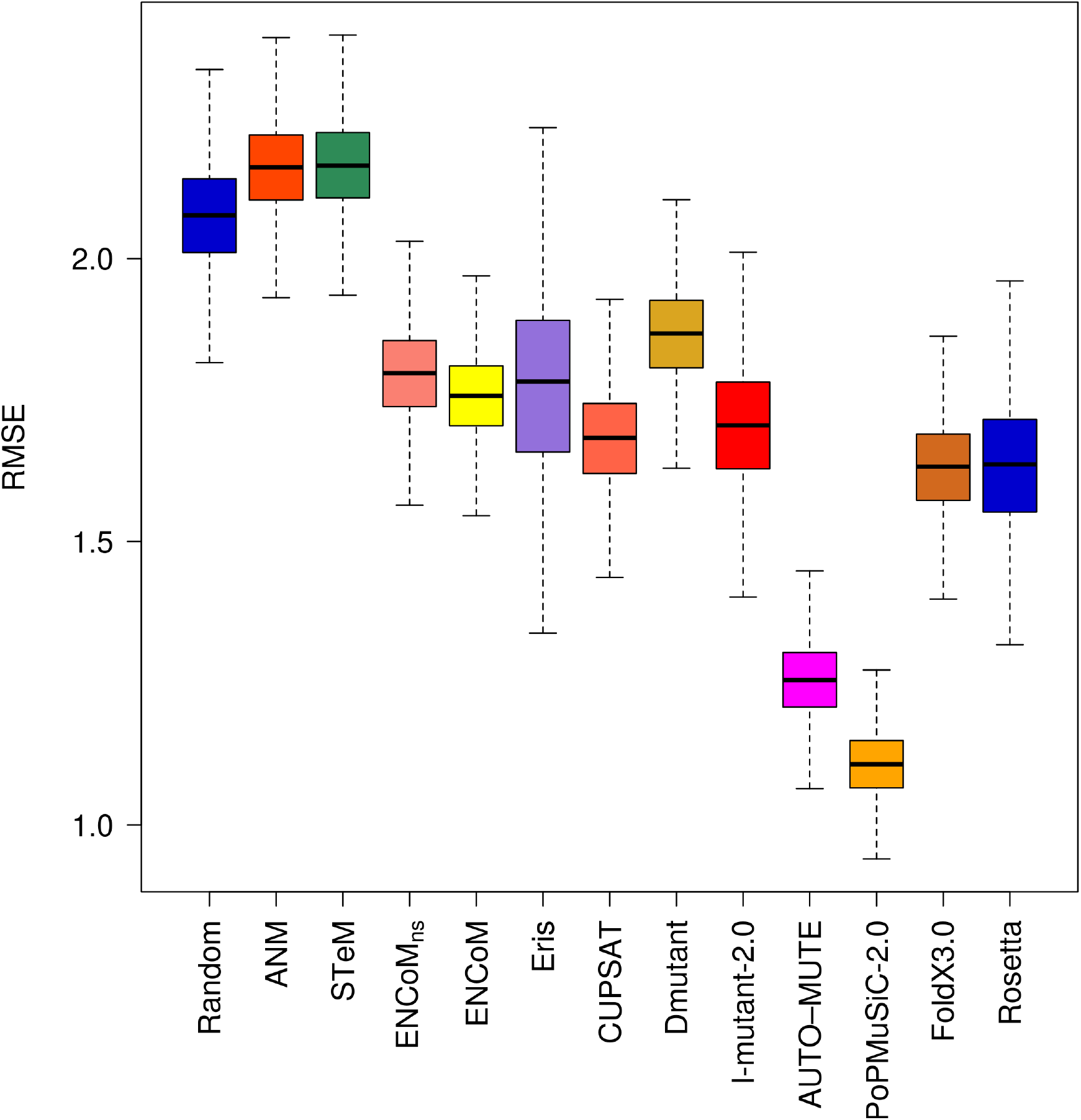
Root Mean Square Error (RMSE) on the prediction of the effect of destabilizing mutations. RMSE of the linear regression through the origin between experimental and predicted variations in free energy variations (ΔΔG). Box plots generated from 10000 resampling bootstrapping iterations on the subset of destabilizing mutations (experimental ΔΔG > 0.5 kcal/mol) in the PoPMuSiC-2.0 dataset (N = 174). The results for destabilizing mutations mirror to a great extent those for the entire dataset given that they represent 57% of the entire dataset.

The subset of 45 stabilizing mutations (Figure 6) gives completely different results as those obtained for destabilizing mutations. AUTO-MUTE, Rosetta, FoldX 3.0 and PoPMuSiC-2.0 that outperformed all of the models on the whole dataset or the destabilizing mutations dataset cannot predict better than the random model. This is also true for CUPSAT, I-Mutant 2.0 and Eris. ENCoM and DMutant are the only models with significantly better than random RMSE values for the prediction of stabilizing mutations.

**Figure 6.**
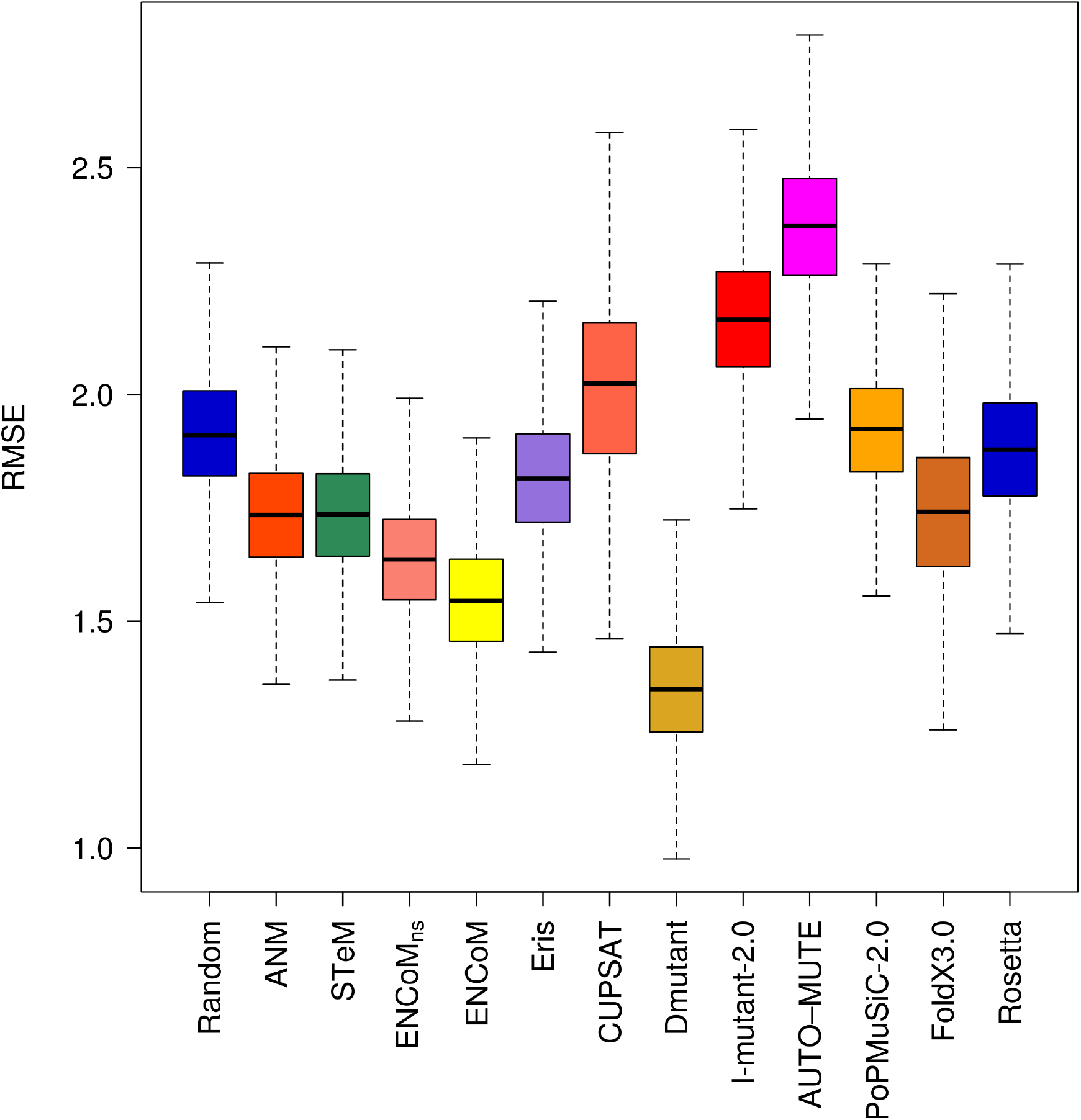
Root Mean Square Error (RMSE) on the prediction of the effect of stabilizing mutations. RMSE of the linear regression through the origin between experimental and predicted variations in free energy variations (ΔΔG). Box plots generated from 10000 resampling bootstrapping iterations on the subset of stabilizing mutations (experimental ΔΔG < −0.5 kcal/mol) in the PoPMuSiC-2.0 dataset (N = 45). With the exception of DMutant and ENCoM, most methods are not substantially better than random and some are substantially worst for stabilizing mutations.

ANM and STeM outperform all models on the neutral mutations (Figure 7). All other models fail to predict neutral mutations any better than random. While the accuracy of ANM and STeM to predict neutral mutations may seem surprising at first, it is in fact an artifact of the methodology. As the wild type or mutated structures are assumed to maintain the same general backbone structure, the eigenvectors/eigenvalues calculated with ANM or STeM will always be extremely similar for wild type and mutant forms. Any differences will arise as a result of small variations in backbone conformation produced by Modeller. As such, ANM and STeM predict almost every mutation as neutral, explaining their high success in this case.

**Figure 7.**
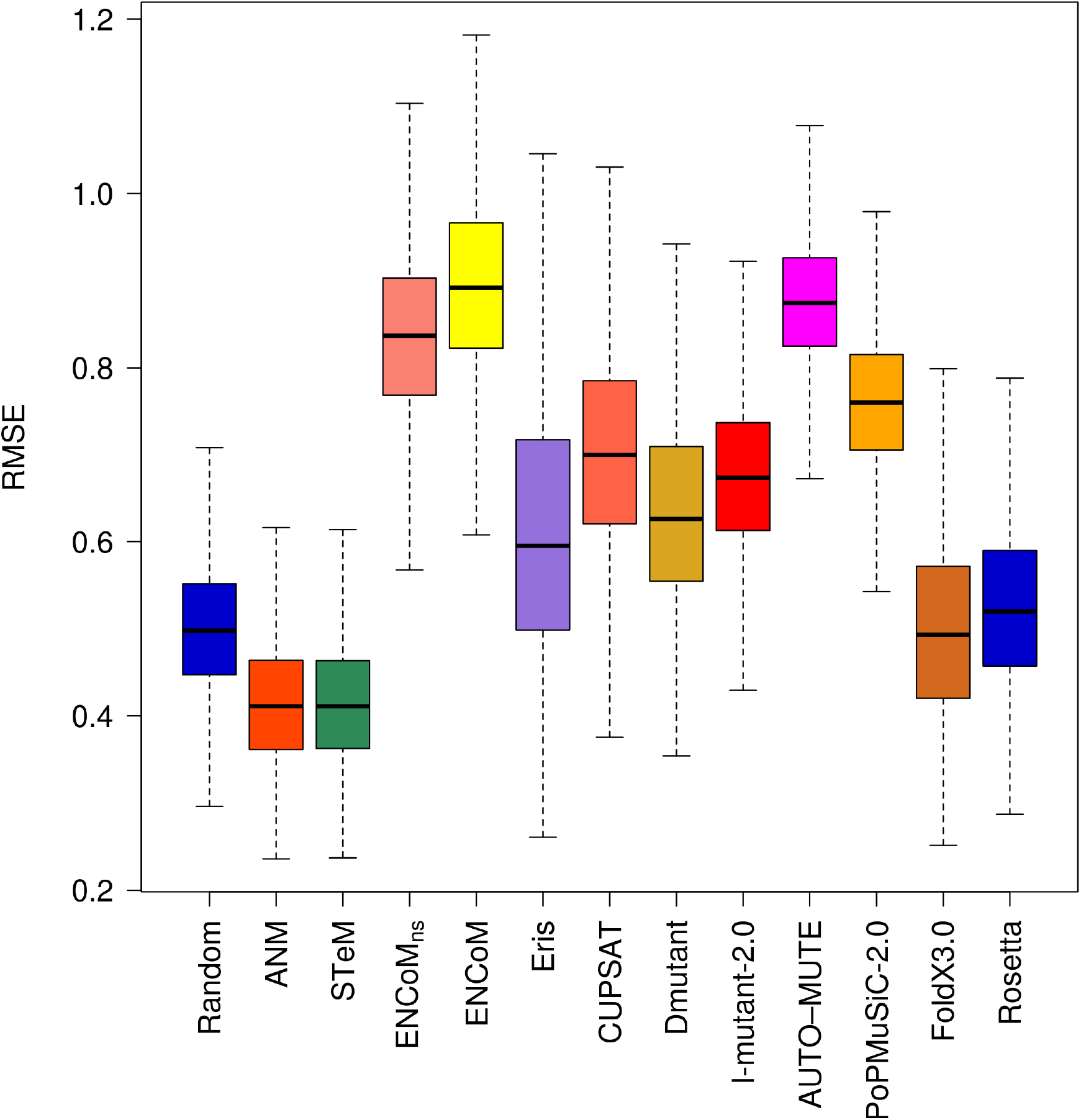
Root Mean Square Error (RMSE) on the prediction of the effect of neutral mutations. RMSE of the linear regression through the origin between experimental and predicted variations in free energy variations (ΔΔG). Box plots generated from 10000 resampling bootstrapping iterations on the subset of neutral mutations (experimental ΔΔG = [-0.5,0.5] kcal/mol) in the PoPMuSiC-2.0 dataset (N = 84). Apart from ANM and STeM that predict most mutations as neutral as an artifact, no method is better than the reshuffled random model at predicting neutral mutations.

At first glance, the comparison of ENCoM_ns_ and ENCoM could suggest that a large part of the effect observed come from a consideration of the total area in contact and not the specific types of amino acids in contact. However, the side chains in contact are already in conformations that minimize unfavorable contacts to the extent that is acceptable in reality (in the experimental structure) or as a result of the energy minimization performed by Modeller for the mutant form given the local environments. The fact that ENCoM is able to improve on ENCoM_ns_ is the actual surprising result and points to the existence of frustration in molecular interactions [64].

Considering that none of the existing models can reasonably predict neutral mutations, the only models that achieve a certain balance in predicting both destabilizing as well as stabilizing mutations better than random are ENCoM and DMutant.

### Self-Consistency in the prediction of the effect of mutations

One basic requirement for a system that predicts the effect of mutations on stability is that it should be self-consistent, both unbiased and with small error with respect to the prediction of the forward or back mutations as reported by Thiltgen *et al.* [65]. The authors built a non-redundant set of 65 pairs of PDB structures containing single mutations (called form A and form B) and utilized different models to predict the effect of each mutation going from the form A to form B and back. From a thermodynamic point of view, the predicted variation in free energy variation should be of the same magnitude for the forward or back mutations, ΔΔG_A->B_ = −ΔΔG_B->A_.

Using the Thiltgen dataset we performed a similar analysis for ENCoM, ENCoM_ns_, ANM, STeM, CUPSAT, DMutant, PoPMuSiC-2.0 and a random model (Gaussian prediction with unitary standard deviation). For the remaining methods (Rosetta, Eris and I-Mutant) we utilize the data provided by Thiltgen. We removed three cases involving prolines as such cases produce backbone alterations. Furthermore, PoPMuSiC-2.0 failed to return results for five cases. The final dataset therefore contains 57 pairs (Supplementary Table IV). The CUPSAT and AUTO-MUTE servers failed to predict 25 and 32 cases respectively. As these failure rates are significant considering the size of the dataset, we prefer to not include these two methods in figures 8 and 9 (the remaining cases appear however in Supplementary Table IV).

**Figure 8.**
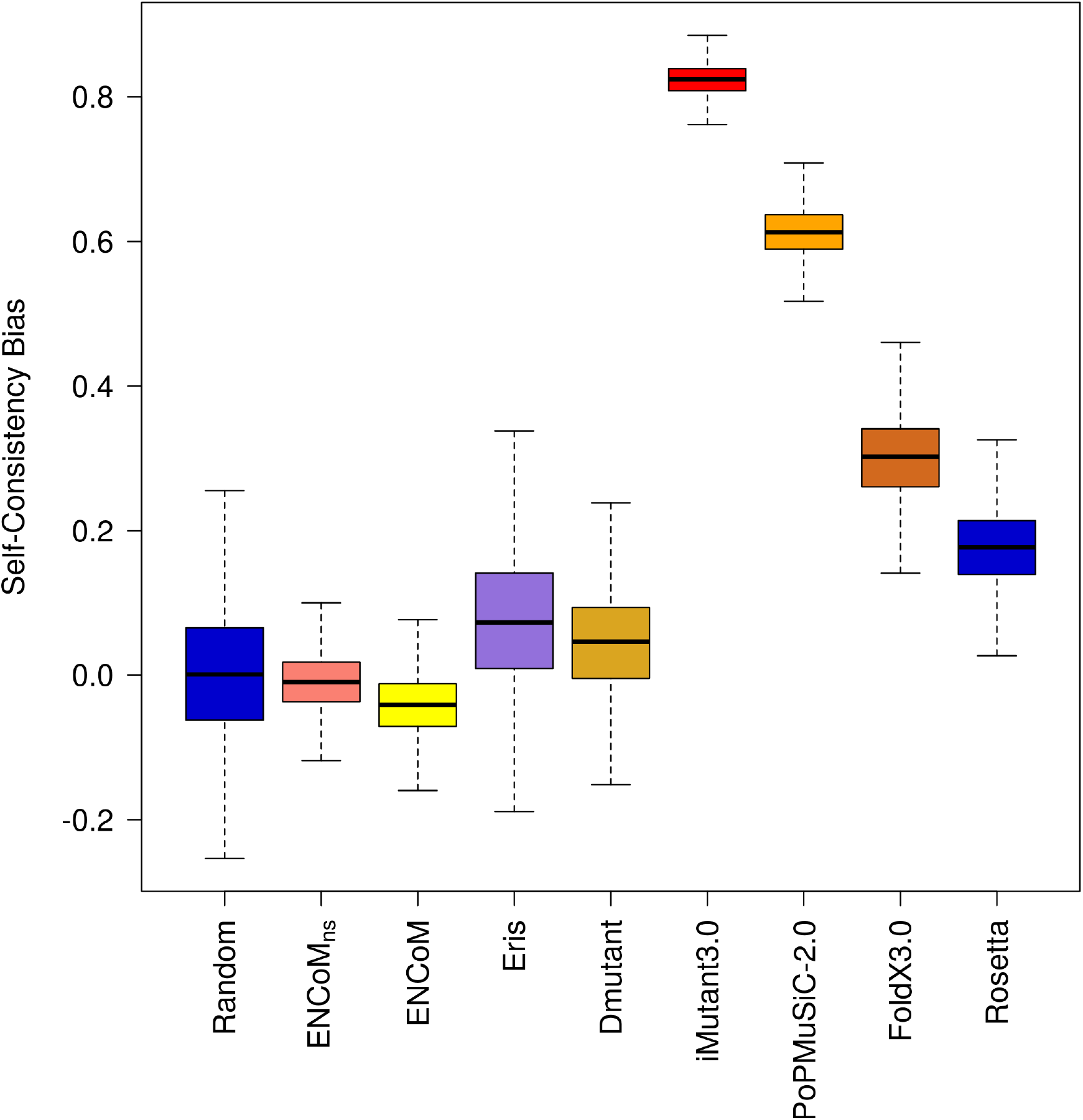
Self-consistency Bias. The bias quantifies the tendency of a method to predict more accurately mutations in one direction than in the opposite. Machine learning based methods in particular show a high bias. ENCoM/ENCoM_ns_ have low bias.

**Figure 9.**
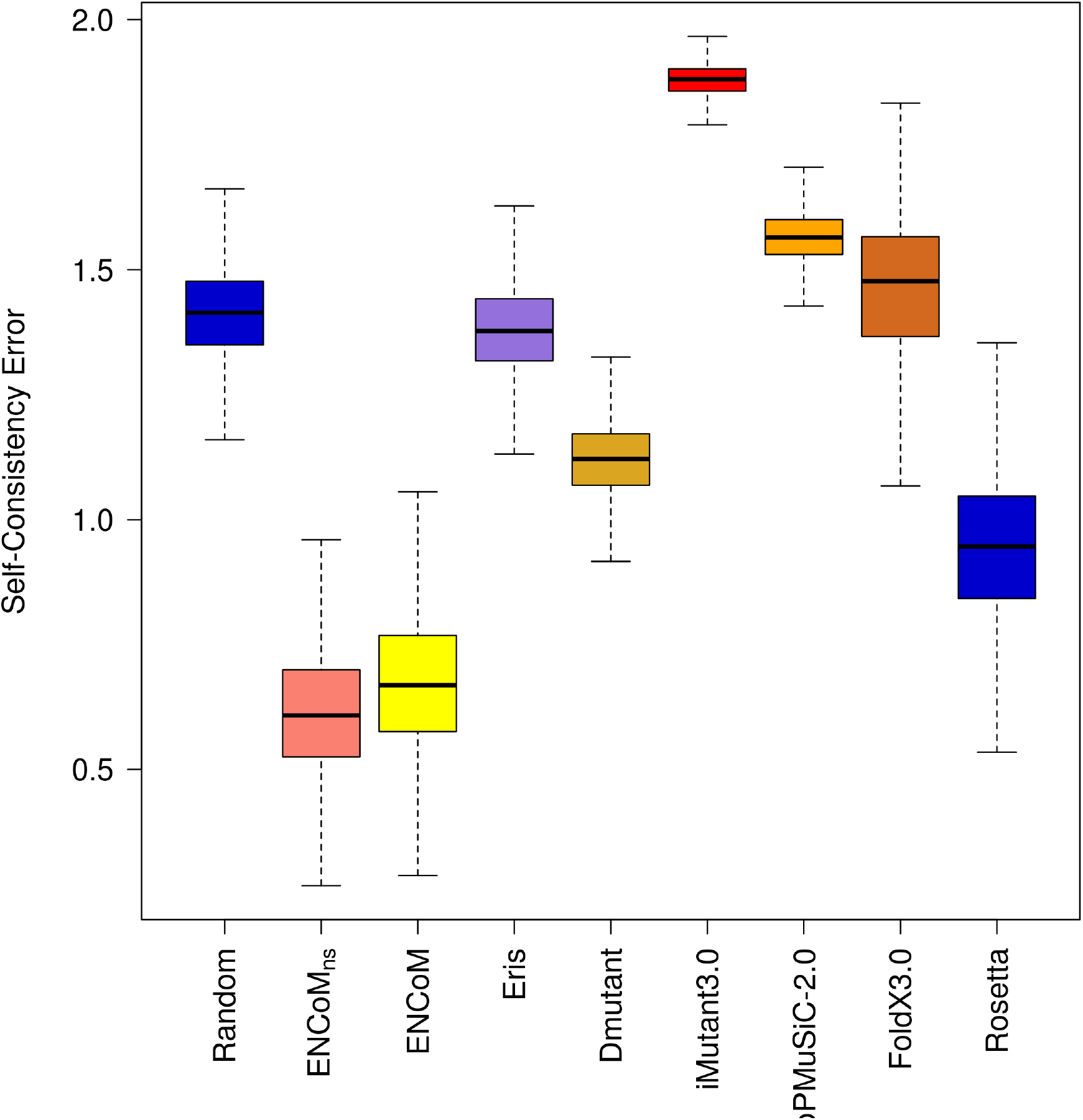
Self-consistency error. The error calculated the magnitude of the biases in the prediction of forward and back mutations. Box plots were generated from 10000 resampling bootstrapping iterations for the 57 proteins pairs in the Thiltgen dataset. ENCoM/ENCoM_ns_ are the methods with lowest self-consistency errors.

The results in figure 8 show that compared to the random model (a positive control in this case), Rosetta and FoldX 3.0 show moderate bias while PoPMuSiC-2.0 and I-Mutant show significant bias. All biased methods are biased toward the prediction of destabilizing mutations (data not shown) in agreement with the results in Figure 3. DMutant, Eris, ENCoM and ENCoM_ns_ are the only models with bias comparable to that of the random model (the positive control in this experiment).

ENCoM, ENCoM_ns_, and to a lesser extent Rosetta and DMutant have lower errors than the random model (Figure 9). All other methods display an error equal or higher than that of the random model. ENCoM and ENCoM_ns_ vastly outperform all the others models in terms of error.

Lastly, STeM and ANM show low and moderate biases respectively and errors equivalent to random (data not shown) but as mentioned, these methods cannot be used for the prediction of mutations (other than neutral mutations as an artifact).

### Prediction of NMR S^2^ order parameter differences

Mutations may not only affect protein stability but also protein function. While experimental data is less abundant, one protein in particular, dihydrofolate reductase (DHFR) from *E. coli*, has been widely used experimentally to understand this relationship [66,67]. Recently, Boher *et al*. [44] have analyzed the effect of the G121 V mutation on protein dynamics in DHFR by NMR spectroscopy. This mutation is located 15Å away from the binding site but reduces enzyme catalysis by 200 fold with negligible effect on protein stability (0.70 kcal/mol). The authors evaluated, among many other parameters, the S² parameter of the folic acid bound form for the wild type and mutated forms and identify the regions where the mutation affects flexibility. We calculated b-factor differences (Equation 4) between the folate-bound wild type (PDB ID 1RX7) and the G121 V mutant (modeled with Modeller) forms of DHFR. We obtain a good agreement (Pearson correlation = 0.61) between our predicted b-factor difference and S^2^ differences (Figure 10). As mentioned earlier, the overall correlation of 0.54 in the prediction of b-factors (Figure 1) appears at least in this case to be sufficient to capture essential functional information.

**Figure 10.**
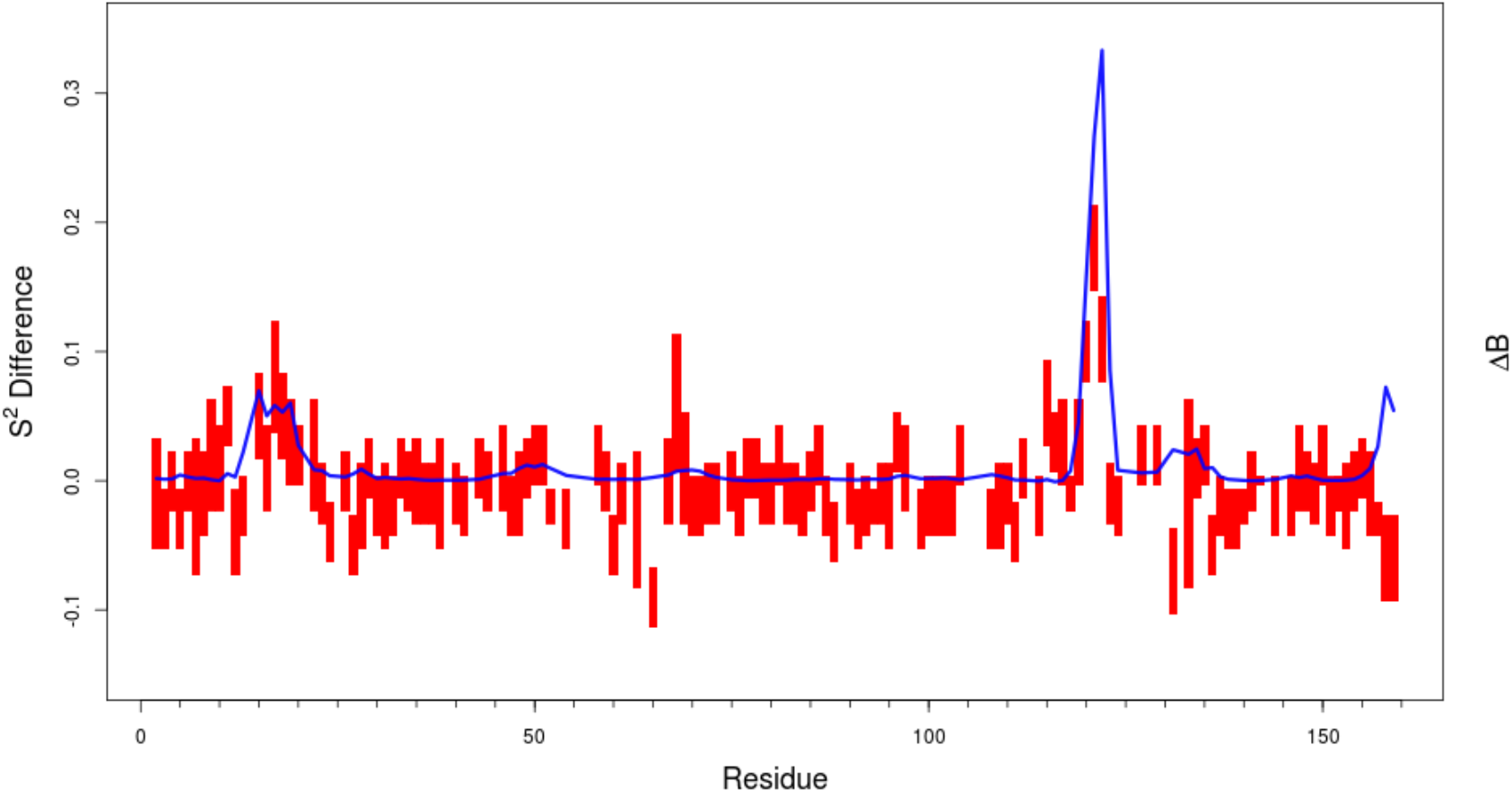
Prediction of NMR S^2^ order parameter differences for the G121 V mutation on DHFR from *E. coli*. Experimental values (red, the bar represents the experimental error) are compared to the inverse normalized predicted b-factors differences, ΔB (Equation 4, blue line) showing a Pearson correlation coefficient of 0.6.

## DISCUSSION

Our results show that a small modification of the long-range interaction term in the potential energy function of STeM had an important positive impact on the model. This small change improves the method in comparison to existing NMA methods in the traditional areas such as the prediction of b-factors and conformational sampling (overlap) where coarse-grained normal mode analysis are applied. More importantly however, it opens an entire new area of application to coarse-grained normal mode analysis methods. Specifically ENCoM is the first coarse-grained normal-mode analysis method that permits to take in consideration the specific sequence of the protein in addition to the geometry. This is introduced through a modification in the long-range interactions to account for types of atoms in contact modulated by their surface in contact. As a validation of the approach we explored the ability of the method to predict the effect of mutations in protein stability. In doing so we created the first entropy-based methodology to predict the effect of mutations on the thermodynamic stability of proteins. This methodology is entirely orthogonal to existing methods that are either machine learning or enthalpy based. Not only the approach is novel but also the method performs extremely favorably compared to other methods when viewed in terms of both error and bias.

As the approach taken in ENCoM is completely different from existing methods for the prediction of the effect of mutations on protein stability, a new opportunity arises to combine ENCoM with enthalpy and machine-learning methods. Unfortunately, we tried to create a naïve method based on linear combinations of the predictions of ENCoM and the different methods presented without success, perhaps due to the large bias characteristic to the different methods.

To assess the relative importance of contact area and the modulation of interactions with atom types, we tested a model that has non-specific atom-type interactions (ENCoM_ns_), this model is atom type insensitive, but is sensitive the orientation of side-chain atoms. While a large fraction of the observed effect can be attributed to surfaces in contact only, ENCoM is consistently better than ENCoM_ns_, particularly at predicting destabilizing mutations where the possibility to accommodate unfavorable interactions is more restricted. We cannot however exclude the effect of the intrinsic difficulty in modeling destabilizing mutations. For stabilizing mutations, the near equivalence of ENCoM and ENCoM_ns_ may be explained in part by the successful energy minimization of the mutated side-chain performed by Modeller. ANM and STeM failed to predict the effect of mutations on the whole dataset. They were not expected to perform well because their respective potentials only take in account the position of alpha carbons (backbone geometry). As such ANM and STeM tend to predict mutations as neutral, explaining their excellent performance onto the neutral subset and failure otherwise. Our results suggest that surfaces in contact are essential in a coarse-grained NMA model to predict the effect of mutation and that the specific interactions between atom types is necessary to get more subtle results, particularly stabilizing mutations.

ENCoM is consistently better than ENCoM_ns_ in the prediction of loop or domain movements irrespective of the dependency of the coupling of this movement to ligand binding or the starting structure (apo or holo form) and both outperform ANM and STeM. Our results corroborate previous work on a mix coarse-grained method adding a atomistic resolution to loops capable of improving the prediction of loop movements [68]. ENCoM performs considerably better than STeM throughout despite having very similar potentials, showing the importance of surfaces in contact in the prediction of movements. There is little difference between ENCoM_ns_ and ENCoM in the prediction of b-factors, but both perform worst than ANM, STeM and GNM. At least in the case of DHFR b-factor differences capture some essential characteristics of the system as calculated by NMR. However, one should be careful in placing too much emphasis on the validation of b-factor predictions using experimental data derived from crystals as these are affected to a great extent by rigid body motions within the crystal [69].

PoPMuSiC-2.0, AUTO-MUTE, FoldX 3.0 and Rosetta perform better than other models in the whole test dataset of mutations. However, the dataset consists of 15% stabilizing mutation, 57% of destabilizing and 28% of neutral mutations. When looking at each subset, machine learning or enthalpy based models failed to predict better than random on the stabilizing mutations subset. Biases in the dataset may have affected the training of machine-learning methods. For example the training set of PoPMuSiC-2.0 contains 2648 mutations in proportions that are similar to those in the testing set with 60%, 29% and 11% destabilizing, neutral and stabilizing mutations respectively. While it is true that most mutations tend to be destabilizing, if one is interested in detecting stabilizing mutations, a method over trained on destabilizing mutations will not meet expectations. Indeed, PoPMuSiC-2.0 and I-mutant the two machine learning based methods, have larger biases and errors than other methods in their predictions. Our method relies on a model structure of the mutant. As the modeling may fail to find the most stable side-chain conformation, it could have a bias toward giving slightly higher energies to the mutant. Notwithstanding this potential bias, ENCoM have the lowest error and bias. This may be a case where less is more as the coarse-grained nature of the method makes it also less sensitive to errors in modeling that may affect enthalpy-based methods to a greater extent. Finally, there is one more advantage in the approach taken in ENCoM. As the network model is a global connected model it considers indirectly the entire protein, while in existing enthalpy or machine-learning methods the effect of a mutation is calculated mostly from a local point of view.

The prediction of the thermodynamic effect of mutations is very important to understand disease-causing mutations as well as in protein engineering. With respect to human diseases, and particularly speaking of cancer mutations, one of the factors that may lead to tumor suppressor or oncogenic mutations is their effect on stability (the authors thank Gaddy Getz from the Broad Institute for first introducing us to this hypothesis). Specifically, destabilizing mutations in tumor suppressor genes or alternatively stabilizing mutations in oncogenes may be driver mutations in cancer. Therefore the prediction of stabilizing mutations may be very important to predict driver mutations in oncogenes. Likewise, in protein engineering, one major goal is that of improving protein stability with the prediction of stabilizing mutations. Such mutations may be useful not as the final goal (for purification or industrial purposes) but also to create a ‘stability buffer’ that permits the introduction of potentially destabilizing additional mutations that may be relevant to create the intended new function.

The work presented here is to our knowledge also the most extensive test of existing methods for the prediction of the effect of mutations in protein stability. The majority of methods tested in the present work fail to predict stabilizing mutations. However, we are aware that the random reshuffled model used may be too stringent given the excessive number of destabilizing mutations in the dataset. The only models that predict stabilizing as well as destabilizing mutations are ENCoM and DMutant, however ENCoM is the only method with low self-consistency bias and error.

While the contribution of side chain entropy to stability is well established [70,71], here we use backbone normal modes to predict stability. As a consequence of the relationship between normal modes and entropy, our results attest to the importance of backbone entropy to stability and increase our understanding of the overall importance of entropy to stability. The strong trend observed on the behavior of different parameters sets with respect to the α_4_ parameter is very interesting. Lower values are associated with better predictions of conformational changes while higher values are associated with better b-factor predictions. One way to rationalize this observation is to consider that higher α_4_ values lead to a rigidification of the structure, adding constraints and restricting overall motion. Likewise, lower α_4_ values remove constraints and thus lead to higher overlap.

We used ENCoM to predict the functional effects of the G121 V mutant of the *E. coli* DHFR compared to NMR data. This position is part of a correlated network of residues that play a role in enzyme catalysis but with little effect on stability. The mutation affects this network by disrupting the movement of residues that are far from the binding site. We can predict the local changes in S^2^ with ENCoM. As these predictions are based on b-factor calculations, this result shows that at least in this case, even with b-factor prediction correlation lower than ANM, STeM and GNM we can detect functionally relevant variations. Clearly, despite the greater performance of GNM, ANM or STeM in the calculation of b-factors, these methods cannot predict b-factor differences as a consequence of mutations, as their predictions are the same for the two forms. While a more extensive study is necessary involving S^2^ NMR parameters, our results serve as an example against relying too heavily on crystallographic b-factors for the evaluation of normal mode analysis methods.

## METHODS

The fundamentals of Normal Mode Analysis (NMA) have been extensively reviewed [72,73]. The key assumption in NMA is that the protein is in an equilibrium state around which fluctuations can be described using a quadratic approximation of the potential via a Taylor series approximation. In equilibrium the force constants are summarized in the Hessian matrix *H* that contains the elements of the partial second derivatives of the terms of the potential with respect to the coordinates. The potential used in ENCoM is similar to that of STeM, a Go-like potential where the closer a conformation 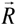 is to the reference (in this case equilibrium) conformation 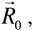 the lower the energy.

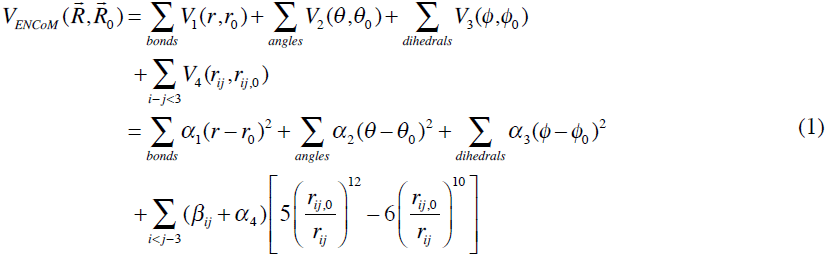

The principal difference between the potential above and that of STeM are the *β_ij_* terms that modulate non-bonded interactions between amino acid pairs according to the surface area in contact. Specifically,

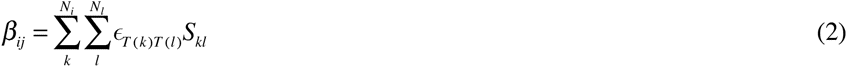

where ∈*_T_*_(*k*_*_)T_*_(_*_l)_* represents a pairwise interaction energy between atom types *T* (*k*) and *T* (*l*) of atom *k* and *l* respectively of amino acids *i* and *j* containing *N_i_* and *N_j_* atoms each. Finally, *S_kl_* represent the surface area in contact between atoms *k* and *l* calculated analytically [45]. We utilize the atom types classification of Sobolev *et al.* [74] containing 8 atom types. A matrix with all the interaction between atom types set at the value 1 is used in the non-specific ENCoM_ns_ model. In Figure 11 we illustrate with the concrete case of a set of 3 amino acids (D11, D45 and R141) in *M. tuberculosis* ribose-5-phosphate isomerase (PDB ID 2VVO), the differences between ENCoM, STeM and ANM in terms of spring strengths associated to different amino acids pairs. D45 is equally distant from R141 and D11 (around 6.0Å) and interacts with R141 but does not with D11. Likewise, D11 does not interact with R141 with a C_α_ distance of 11.6Å. ANM assigns equal strength to all three pairwise C_α_ springs (as their distances fall within the 18.0Å threshold). STeM assigns equal spring strengths to the D11-D45 and D45-R141 pairs. Among the three methods, ENCoM is the only one to properly assign an extremely weak strength to the D11-R141 pair, a still weak but slightly stronger strength to the D11-D45 pair (due to their closer distance) and a very strong strength to the spring representing the D45-R141 interactions.

**Figure 11.**
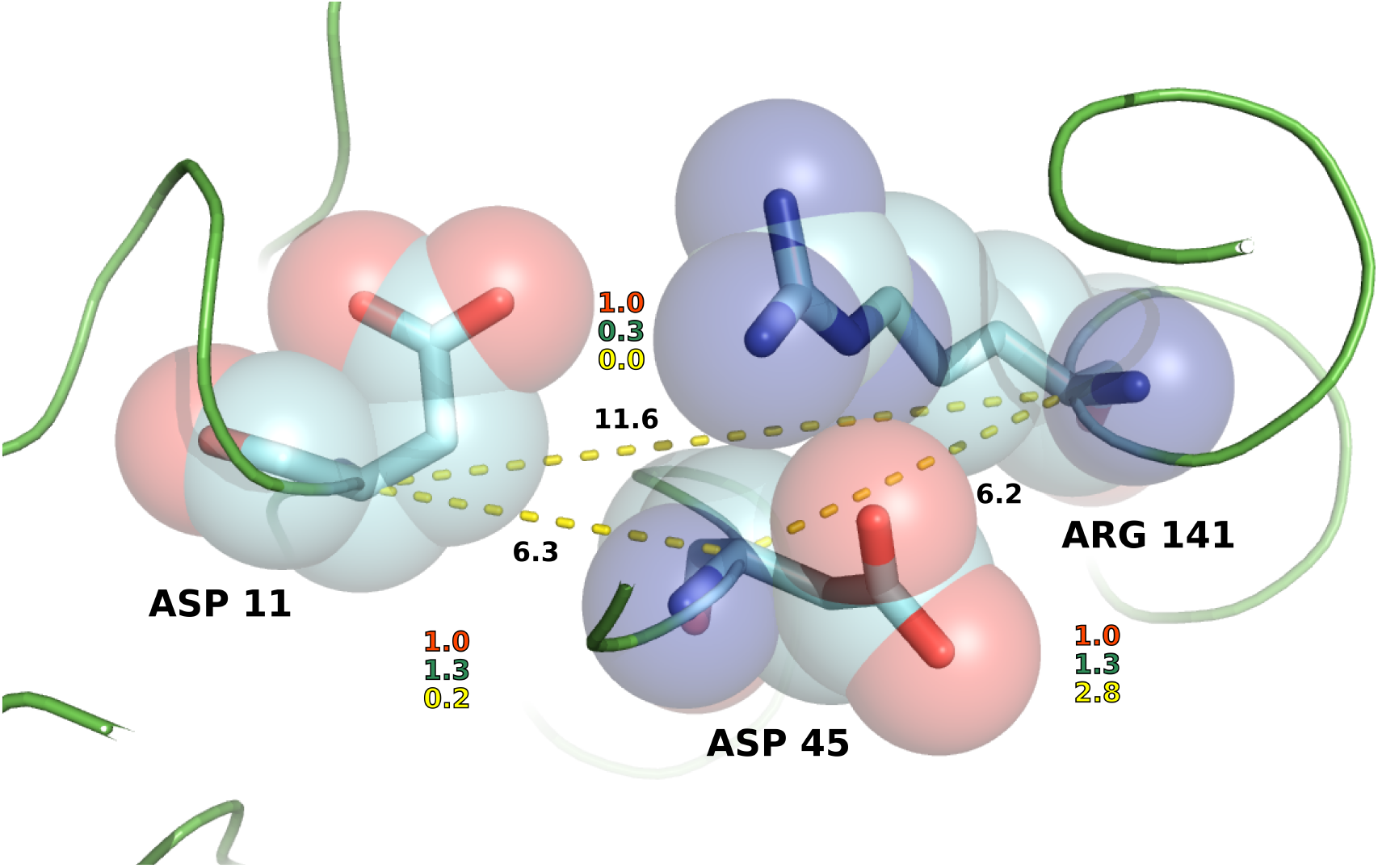
Illustration of the representation of inter-residue interactions by the different NMA methods. The figure shows three amino acids (D11, D45 and R141) from the *M. tuberculosis* ribose-5-phospate isomerase (PDB ID = 2VVO). The distances between alpha carbons are shown with the yellow dotted lines and labeled in black. Interaction strengths relative to ANM (in red) are shown for STeM (green) and ENCoM (yellow). ANM treats all pairs as equal while STeM treat equally D11 and R141 with respect to D45 even though the interaction between D45 and R141 is much stronger by virtue of side-chain interactions, as correctly described in ENCoM.

The hessian matrix can be decomposed into eigenvectors 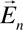 and their associated eigenvalues *γ_n_*. For a system with *N* amino acids (each represented by one node in the elastic network), there are 3*N* eigenvectors. Each eigenvector describes a mode of vibration in the resonance frequency defined by the corresponding eigenvalue of all nodes, in other words the simultaneous movement in distinct individual directions for each node (C_α_ atoms in this case). The first 6 eigenvectors represent rigid body translations and rotations of the entire system. The remaining eigenvectors represent internal motions. The eigenvectors associated to lower eigenvalues (lower modes) represent more global or cooperative movements while the eigenvectors associated to higher eigenvalues (fastest modes) represent more local movements. Any conformation of the protein can be described by a linear combination of different amplitudes 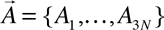 of eigenvectors 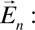

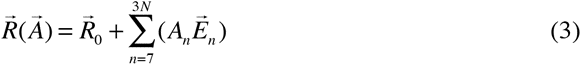

The source code for ENCoM is freely available at http://bcb.med.usherbrooke.ca/encom.

### Parameterization of ENCoM

In order to obtain a set of *α* parameters to be used with ENCoM we performed a sparse exhaustive integer search of the logarithm of *α* parameters with *α_i_* = [10^−4^, 10^8^] for *i* = [1, 4] to maximize the prediction ability of the algorithm in terms of overlap and prediction of mutations. In other words, we searched all combinations of 13 distinct relative orders of magnitude for the set of 4 parameters *α*. For each parameter set, we calculated the bootstrapped median RMSE (see below) Z-score sum for the prediction of stabilizing and destabilizing mutations, 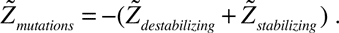 Keeping in mind that lower RMSE values represent better predictions, the 2000 parameter sets (out of 28561 combinations) with highest 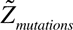 were then used to calculate Z-scores for overlap in domain and loop movements. As our goal is to obtain a parameter set that combines low RMSE and high overlap, we ranked the 1000 parameter sets according to 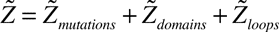. The parameter set with highest 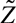 is (*α*_1_, *α*_2_, *α*_3_, *α*_4_) = (10^2^, 10^4^, 10^4^, 10^−2^) (solid black line in Figure 12). The optimization of the bootstrapped median is equivalent to a training procedure with leave-many-out testing.

The exploration of parameter space shows that there is a clear trade-off between the prediction of mutations (low RMSE), conformational sampling (high overlap) and b-factors (high correlations). Parameter sets that improve the prediction of b-factors are invariably associated with poor conformational changes (low overlap) associated to both domain and loop movements and variable RMSE for the prediction of mutations (red lines in Figure 12). On the other hand, parameter sets that predict poorly b-factors, perform better in the prediction of conformational changes and the effect of mutations (blue lines in Figure 12). The *α* parameters used in STeM, (*α*_1_, *α*_2_, *α*_3_, *α*_4)_ = (360, 72, 9.9, 3.6) are arbitrary, taken without modifications from a previous study focusing on folding [75]. As expected, this set of parameters can be considerably improved upon as can be observed in Figure 12 (dashed line).

**Figure 12.**
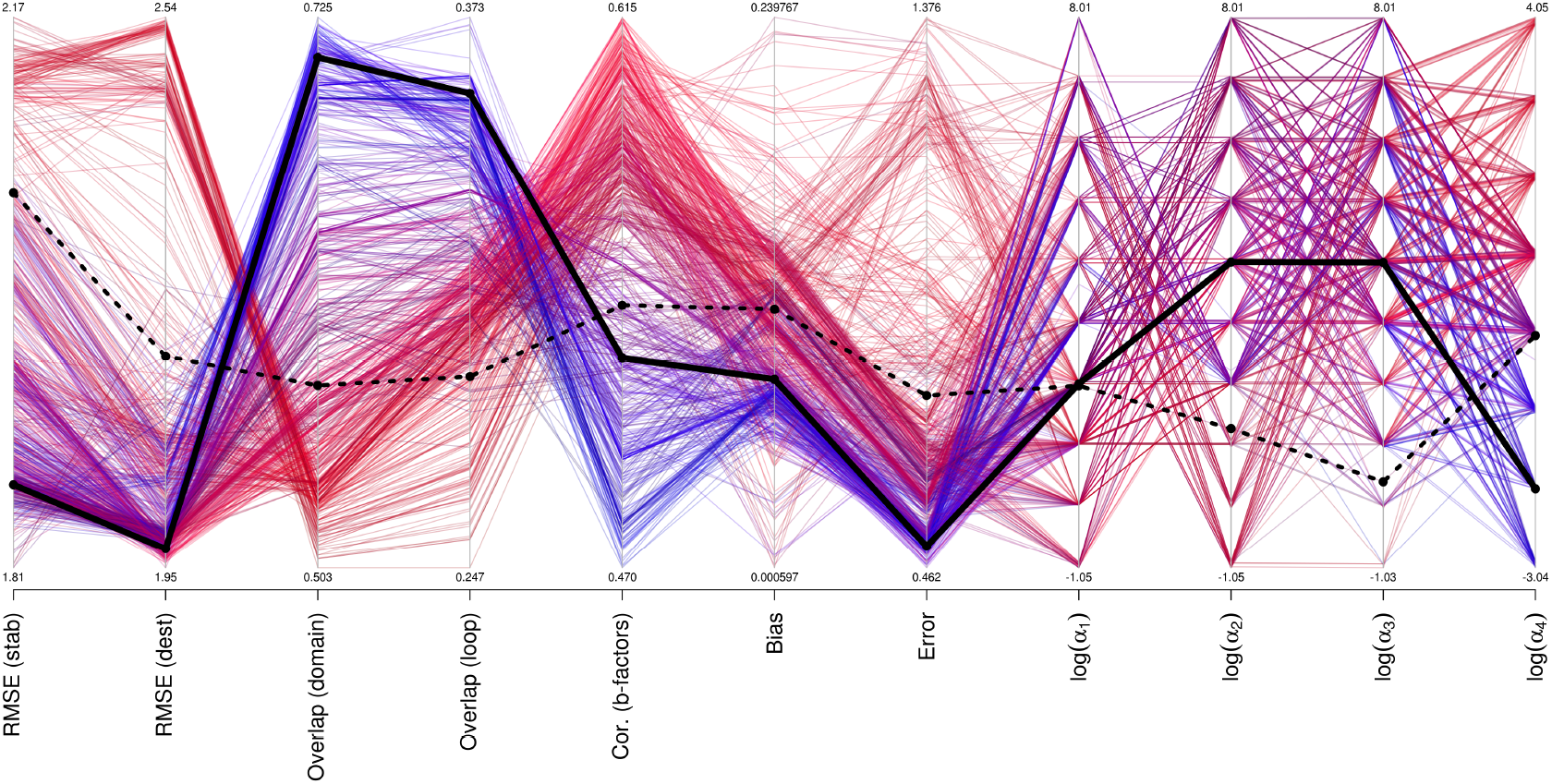
Performance of different parameter sets on the prediction of mutations, b-factors and motions. We present as a parallel plot the bootstrapped median RMSE for stabilizing and destabilizing mutation, average best overlap for domains and loop movements as well as self-consistency bias and errors. In the right-most four columns with include the logarithm of the 4 alpha variables. Different parameter sets are colored based on b-factors correlations (red gradient) or domain movement overlaps (blue gradient). The black line represent the specific set of parameters used in ENCoM while the dashed line represents the values for ENCoM using the set of alpha parameters employed in STeM. There is a dichotomy in parameter space such that most sets of parameters are either good at predicting b-factors or overlap and mutations.

The four right-most variables in the parallel coordinates plot in Figure 12 show the logarithm of the α parameters for each parameter set. Either class of parameter sets, better for b-factors (in red) or better for overlap/RMSE (in blue) come about from widely diverging values for each parameter across several orders of magnitude. There are however some patterns. Most notably for α_4_, where there is an almost perfect separation of parameter sets around α_4_ = 1. Interestingly, higher values of α_1_ and α_2_, associated with stronger constraints on distances and angles tend also to be associated to better overlap values. While it is likely that a better-performing set of parameters can be found, the wide variation of values across many orders of magnitude show that within certain limits, the method is robust with respect to the choice of parameters. This result justifies the sparse search employed.

### Bootstrapping

Bootstrapping is a simple and general statistical technique to estimate standard errors, p-values, and other quantities associated with finite samples of unknown distributions. In particular, bootstrapping help mitigate the effect of outliers and offers better estimates in small samples. Bootstrapping is a process by which the replicates (here 10000 replicates) of the sample points are stochastically generated (with repetitions) and used to measure statistical quantities. In particular, bootstrapping allows the quantification of error of the mean [76–80]. Explained in simple terms, two extreme bootstrapping samples would be one in which the estimations of the real distribution of values is entirely made of replicates of the best case and another entirely of the worst case. Some more realistic combination of cases in fact better describes the real distribution. Thus, bootstrapping, while still affected by any biases present in the sample of cases, helps alleviate them to some extent.

### Predicted b-factors

One of the most common types of experimental data used to validate normal mode models is the calculation of predicted b-factors and their correlation to experimentally determined b-factors. B-factors measure how much each atom oscillates around its equilibrium position [6]. Predicted b-factors are calculated as previously described [35]. Namely, for a given C_α_ node (*i*), one calculates the sum over all eigenvectors representing internal movements (n = 7 to 3N) of the sum of the squared *i*^th^ component of each eigenvector in the spatial coordinates *x,y* and *z* normalized by the corresponding eigenvalues:

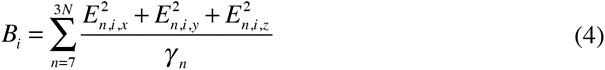

We calculate the Pearson correlation between predicted and experimental b-factors for each protein and average random samples according to the bootstrapping protocol described above.

### Overlap

The overlap is a measure that quantifies the similarity between the direction of movements described by eigenvectors calculated from a starting structure and the variations in coordinates observed between that conformation and a target conformation [49,50]. In other words, the goal of overlap is to quantify to what extent movements based on particular eigenvectors can describe another conformation. The overlap between the n^th^ mode, *O_n_*, described by the eigenvector 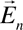 is given by

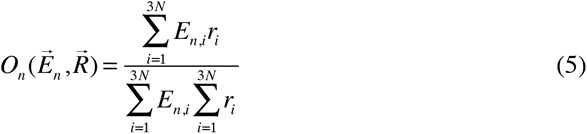

where 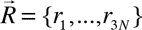 represent the vector of displacements of coordinates between the starting and target conformations. The larger the overlap, the closer one can get to the target conformation from the starting one though the movements defined entirely and by the *n*^th^ eigenvector. We calculate the best overlap among the first 10 slowest modes representing internal motions.

### Evaluating the effect of mutations on dynamics

Insofar as the simplified elastic network model captures essential characteristics of the dynamics of proteins around their equilibrium structures, the eigenvalues obtained from the normal mode analysis can be directly used to define entropy differences around equilibrium. Following earlier work [51,52], the vibrational entropy difference between two conformations in terms of their respective sets of eigenvalues is given by:

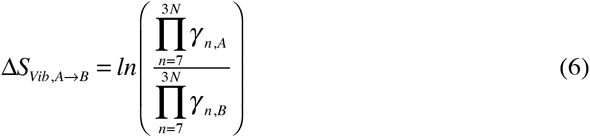

In the present work the enthalpic contributions to the free energy are completely ignored. Therefore, in the present work we directly compare experimental values of ΔG to predicted ΔS values. In order to use the same nomenclature as the existing published methods, we utilize ΔΔG to calculate the variation of free energy variation as a measure of conferred stability of a mutation.

### Root mean square error

A linear regression going through the origin is build between predicted ΔΔG and experimental ΔΔG values to evaluate the prediction ability of the different models. The use of this type of regression is justified by the fact that a comparison of a protein to itself (in the absence of any mutation) should not have any impact on the energy of the model and the model should always predict an experimental variation of zero. However, a linear regression that is not going through the origin would predict a value different from zero equal to the intercept term. In other words, the effect of two consecutive mutations, going from the wild type to a mutated form back to the wild type form (WT->M->WT) would not end with the expected net null change.

The accuracy of the different methods was evaluated using a bootstrapped average root mean square error of a linear regression going through the origin between the predicted and experimental values. We refer to this as RMSE for short and use it to describe the strength of the relationship between experimental and predicted data.

### Self-consistency bias and error in the prediction of the effect of mutations

If one was to plot the predicted energies variation of ΔΔG_A->B_ versus ΔΔG_B->A_ and trace a line y = −x, the bias would represent a tendency of a model to have points not equally distributed above or below that line while the error would represent how far away a point is from this line. In other words, considering a dataset of forward and back predictions, the error is a measure of how the predicted ΔΔG differ and the bias how skewed the predictions are towards the forward or back predictions [65]. A perfect model, both self-consistent and unbiased, would have all the points in the line. Statistically, the measures of bias and error are positively correlated. The higher the error for a particular method, the higher the chance of bias.

## ACKNOWLEDGEMENTS

The authors would like to thank Profs. Pierre Lavigne and Jean-Guy LeHoux from the department of Biochemistry, Université de Sherbrooke for useful discussions throughout the development of the method. VF was partially funded by the Canadian Institutes of Health Research Grant MT-10983 (to Pierre Lavigne and Jean-Guy LeHoux), PROTEO (the Québec network for research on protein function, structure and engineering), Fonds de Recherche du Québec en Santé (FRQ-S) and Nature et Technologie (FRQ-NT). R.J.N. is part of Centre de Recherche Clinique Étienne-Le Bel, a member of the Institute of Pharmacology of Sherbrooke, PROTEO and GRASP (Groupe de Recherche Axé sur la Structure des Protéines) and is the recipient of a Junior Researcher I fellowship from FRQ-S. The funders had no role in study design, data collection and analysis, decision to publish, or preparation of the manuscript.

